# Response of *Asplenium nidus* to Drought Stress and Identification of Drought Tolerance Genes

**DOI:** 10.1101/2020.02.06.936773

**Authors:** Jingwen Liang, Wengang Yu, Peng Wang, Xudong Yu, Zeping Cai, Shitao Xu

## Abstract

To explore how *Asplenium nidus* responds to drought stress, and to mine drought-responsive genes for *Asplenium nidus*, we conducted a pot experiment in the IOT smart greenhouse. We measured a series of physiological and biochemical indexes for drought-treated plants, and analyzed the expression of drought-responsive genes in *Asplenium nidus* by RT-qPCR. The results show that the *Asplenium nidus* is a species with drought tolerance. *Asplenium nidus* inhibits plant photosynthesis and reduces life activities by limiting stomatal opening to adapt to drought. To resist drought stress, the *Asplenium nidus* changes the osmotic potential by increasing the proline content to maintain normal metabolic activities and prevents the damage of active oxygen by increasing the enzyme activities of SOD and POD. Based on the analysis of the relative expression levels of genes, *AVP1-2* and *AVP1-4* may be drought-resistant genes in *Asplenium nidus*. This study lays the foundation for in-depth research on drought tolerance mechanisms and drought-resistant gene mining of *Asplenium nidus.*

## Introduction

*Asplenium nidus*, also known as bird’s nest Fern, is a species under the Aspleniaceae family. It is epiphytic on the trunk or rock under the forest. It is a specie of medicinal, edible, ornamental, and purifying epiphytic fern. Besides basic nutrients, the young leaves contain 18 kinds of amino acids, rich mineral elements, and abundant trace elements like Zn, Mu, Gu, etc. (XU Shitao et al., 2012) As a result, it can not only be used as wild vegetables, but also contain medicinal value. For example, it can strengthen muscles and bones, promote blood circulation, remove blood stasis, promote bone cell growth, accelerate fracture healing,(CHEN Qing-yu et al., 2011) cure fever, chest pain, and cough(Benjamin A et al., 2007). In addition, the bird’s nest fern is a representative of the epiphytes in the rainforest with a beautiful shape and a unique nest. The bird’s nest fern can be used to beautify the indoor environment and to absorb harmful gases such as formaldehyde and nicotine(LI JUAN et al., 2009). Therefore, it is popular among people as a natural air purifier.

The application function of Bird’s Nest Fern has gradually been observed, and more and more researchers have began to study its ecological function in the rainforest. Through these investigations and studies, people have realized that Bird’s Nest Fern plays a non-negligible role in tropical rain forest ecosystems. Bird’s Nest Fern has the function of enriching and maintaining forest biodiversity and genetic diversity. It can use precipitation and litter to form perched soil through the processing of invertebrates and microorganisms. High-level soil can not only serve nutrients for the bird’s nest fern, but also share nutrients and water through rainwater, host, and other anthropogenic plants (Fayle et al., 2008). In addition, the perched soil provide other animated plants and microorganisms with habitat, and provide habitat and food for bird’s(DANG Hui et al., 2011). Bird’s Nest Fern has the function of regulating water and cycling nutrient in forest ecosystems (Tom M. Fayle et al., 2009). Its leaves have special water-holding properties (Ainuddin N A et al., 2009). The aerial roots in the nest base can effectively absorb the moisture in the air, and provide moisture for the surrounding microenvironment through root growth during drought period(Weathers K C, 1999). Due to its large nest base, it can collect litter and convert it into high soil to retain water and provide nutrients (TURTON, 2007).

Although the current research interest in bird’s nest ferns is gradually increasing, it is still in its infancy, the understanding of bird’s nest ferns has not been deepened, the ecological environment is destroyed, the survival of bird’s nest ferns has been threatened, and it is urgent to understand its drought tolerance mechanism. When a plant is subjected to drought stress, water redistribution in various parts, injury of membranes and nuclei, inhibition of growth of leaves and roots, weakened photosynthesis, and excessive production of reactive oxygen species will occur. Some drought-resistant plants will undergo a series of changes to resist drought, such as adjusting their own life cycle to avoid drought, controlling stomatal loss through stomatal adjustment, and changing the concentration of solutes in cells through osmotic adjustment to cause cell osmotic potential changes to maintain more normal metabolic activities in plants, or to activate genes that are adapted to adversity and express new proteins to resist drought stress. *AVP1* gene, or proton pyrophosphatase gene, which is synthesized by proton pyrophosphatase gene, is a kind of hydrolytic enzyme with pyrophosphate as substrate and a kind of proton pump, and exists on the tonoplast or cell membrane of plants, a small number of algae and photosynthetic bacteria, can be coupled with H^+^ transmembrane transport, and works with H^+^-ATPase to pump H^+^ in the cytoplasm into vacuoles. It works by maintaining a certain H^+^ gradient between vacuole and cytoplasm, and providing power for the secondary transport of ions and other solutes (amino acids, sugars, etc.) (Hsu et al., 2018). Studies have shown that overexpression of vacuolar proton pyrophosphatase gene can improve salt tolerance and drought resistance in *Arabidopsis thaliana* (Liu Liang et al., 2011), as well as conferring salt tolerance and drought resistance in crops such as tomato (Shimna Bhaskaran et al., 2011), cotton (Hong Zhang et al., 2011), apple(Dong et al., 2011).

After comprehensive consideration of the above contents, bird’s nest fern was treated with different degrees of drought in this experiment. The leaf growth and stomata under the leaf epidermis were observed, and leaf water content and photosynthetic index were measured. The closely related osmotic regulation compound content and enzyme activity were determined. We also analyzed the expression of proton pyrophosphatase gene (*AVP1*) related to drought tolerance to explore the response mechanism of bird’s nest fern to drought stress. This study will provide a theoretical basis for the further study of drought tolerance mechanism and drought tolerance gene evolution mechanism of bird’s nest fern, and offer useful reference for ornamental cultivation and wild bird’s nest fern protection.

## Materials and methods

### Experiment design

This experiment was performed in the IOT smart greenhouse on the Danzhou campus of Hainan University. The experimental materials were 18 plants of *Asplenium nidus* with good growth and uniformity, which were cultivated in a soft plastic nutrition bowl (8cm × 12cm). The culture substrate is coconut bran, husks, peanut husks, the ratio is 1: 1: 1. These plants were divided into three groups of six plants each. After two days of normal daily watering and incubation, proceed as follows (the watering volume is just after water seeps from the basin, and the watering time is fixed at 5 pm every day):

Group 1: control group(CK), was watered every day in the control group.

Group 2: mild drought (T1), irrigated once every two weeks.

Group 3: severe drought (T2), no watering all the time.

All the leaves of T2 group wilted and yellowed (the 6th week) and then rehydrated for two weeks.

### Determination of growth parameters

In this experiment, we measured leaf length and leaf width of the three groups of bird’s nest fern. The measurement time of each indicator is fixed at 9 am. The leaf length and width are measured once a week and measured with a ruler. Each plant selects a growing leaf to track and measure until the leaf stops growing. Calculate weekly increments separately.

### Determination of physiological parameters

In this experiment, we measured five physiological parameters: relative leaf water content(RLWC), the maximum net photosynthetic rate(*P*_*max*_), transpiration rate (*T*_*r*_), stomatal conductance (*g*_*s*_), and intercellular CO_2_ concentration (*C*_*i*_) the three groups of bird’s nest fern.

The RLWC of leaves is measured every two weeks. We randomly selected three plants per treatment, and each plant took a mature and healthy leaf for measurement. The relative water content is measured by measuring the fresh weight (FW) of leaves immediately after removing it, and then immersing leaves in distilled water for 3-4 hours. When the constant weight is reached, the weight of the leaves is saturated weight (TW). Then put leaves into an oven at 120 ° C for 30 minutes and bake it to a constant weight at 90 ° C. At this time, the weight of leaves is dry weight (DW), and the relative leaf water content of the blade is calculated as:

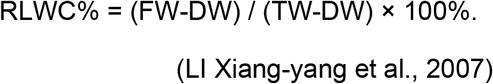

*P*_*max*_, *T*_*r*_, *g*_*s*_, and *C*_*i*_ were measured once a week and measured using a Li-6400 portable photosynthesis instrument (LiCor, Lincoln, NE, USA). Using a red-blue light leaf chamber, the light intensity was set to 1200µmol m^-2^ s^-1^. We select a healthy leaf in each bird’s nest fern, and each leaf is measured once at the tip, middle and base of the leaf.

### Determination of Biochemical parameters

We randomly selected three bird’s nest ferns from each treatment every two weeks. One leaf of each plant was randomly selected and used to determine the content of proline and betaine, and the enzyme activities of superoxide dismutase, peroxidase and catalase. These five parameters were measured using a kit from *Coming Bio Co., Ltd* and Microplate reader(Thermo Multiskan FC, USA).

### Bioinformatics Analysis

Entering the protein sequence of *AVP1* into the PfamA database for searching with standard parameters (E-value less than 1e-5, the length of the domain is more than 10% of the full length of the gene), the information of the amino acid site of the conserved domain of the *AVP1* gene can be obtained.(Finn et al., 2016)

The sequences containing the same conserved domain as *AVP1* in the protein model database of *Asplenium nidus* and *Arabidopsis thaliana* (model plant) were mined by HMMER and BLAST programs with default parameters(E-value=1e-5), respectively, and the results of the two programs were screened and analyzed together (Wang Peng et al., 2018a). The sequences of bird nest fern and *Arabidopsis thaliana* with the same conserved domain as *AVP1* were obtained (Johnson et al., 2008).

We used ProbCons for sequence alignment, compared the protein sequences with the same domain as *AVP1*, which were previously mined in the target species, and then used the PhyML to construct a phylogenetic tree of these protein sequences(Wang Peng et al., 2017), and performed 1,000 bootstrap analysis (Guindon et al., 2009), using Figtree to display the evolutionary tree construction results(Wang Peng et al., 2018b).

### Analysis of gene expression

We randomly selected three bird’s nest ferns from each treatment every two weeks, and randomly selected one leaf from each plant for RNA extraction. We used TIANGEN’s polysaccharide polyphenol plant total RNA extraction kit to extract total RNA from the leaves of Bird’s Nest Fern.

We used the TranScript One-Step gDNA Removal and cDNA Synthesis SuperMix Reverse Transcription Kit to synthesize cDNA templates. Configure the reverse transcription system according to 50ng-5μg Total RNA, 1μl AnchoredOligo (dT) 18Primer (0.5μg/μl), 1μl 2xTS Reaction Mix, 1μl TransScript® RT Enzyme Mix. Reverse transcription was performed at 42°C for 30min, and then heated at 85°C for 5s. Then the cDNA was stored at −20°C.

Primer5 and Oligo7 software were used to design primers for RT-qPCR. The relative expression of genes was detected by a real-time PCR instrument(*ThermoPikoReal,* America). Configure the reverse transcription system according to 1μl cDNA, 0.3μl PCR Forward Primer(10μm), 0.3μl PCR Reverse Primer(10μm), 5μl TransStart Tip Green qPCR SurperMix and 3.4μl ddH_2_O. The relative gene expression was analyzed by 2^∼△△CT^ method.

### Statistical Analysis

All the data were made statistics and calculations with Microsoft Excel 2019. The data in the chart is the mean ± Standard error. The data were statistically analyzed by the Two-way ANOVA program in IBM SPSS Statistics 23.0. Comparisons of results are marked on the chart using the alphabet method by Adobe Illustrator CS5. We use R to analyze the correlation of photosynthetic indicators in each period of T1 and T2. R Studio was used to analyze the correlation of photosynthetic indexes in T1 and T2. The data in the chart is the Pearson Correlation Coefficient and Significance.

## Results

### Measurement results of growth parameters

The increments of leaf length and leaf width in CK group only fluctuated slightly, and there was no significant difference in the increments of leaf length and leaf width of control group during different period (Fig. 2). The leaf length and leaf width of T1 group changed with watering frequency (watering once every two weeks), and could return to normal growth immediately after rehydration (6th week). The plants in T2 group was not watered all the time, and the leaf length and width stopped growing from the 2nd week. Although the growth resumed immediately after rehydration, the growth rate was slower than that of CK group and T1 group. Although it did not return to normal growth rate after two weeks of rehydration, it continued to increase.

**Fig. 1.**
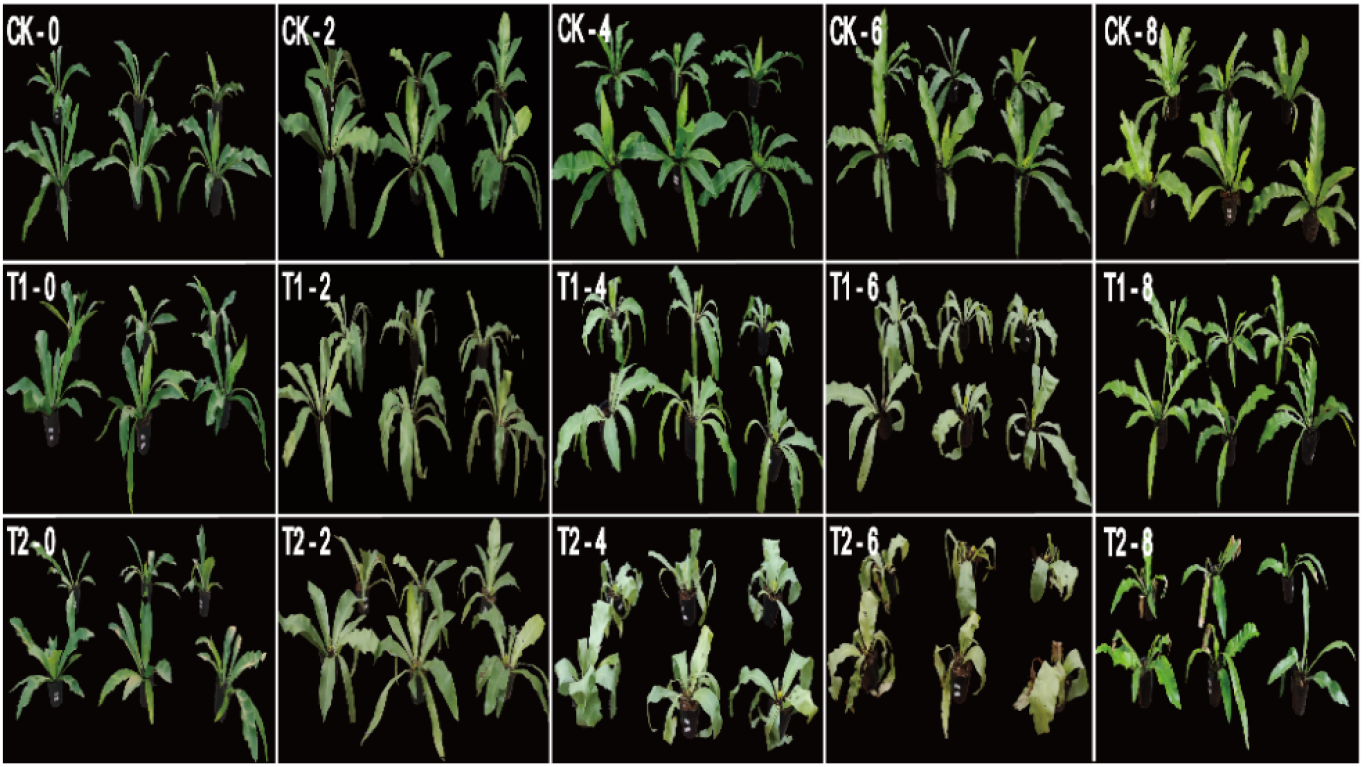
Morphological changes of three groups (CK, T1, T2) of bird’s nest fern with treatment time(0-8week). The 6th week is rehydration time.

**Fig. 2.**
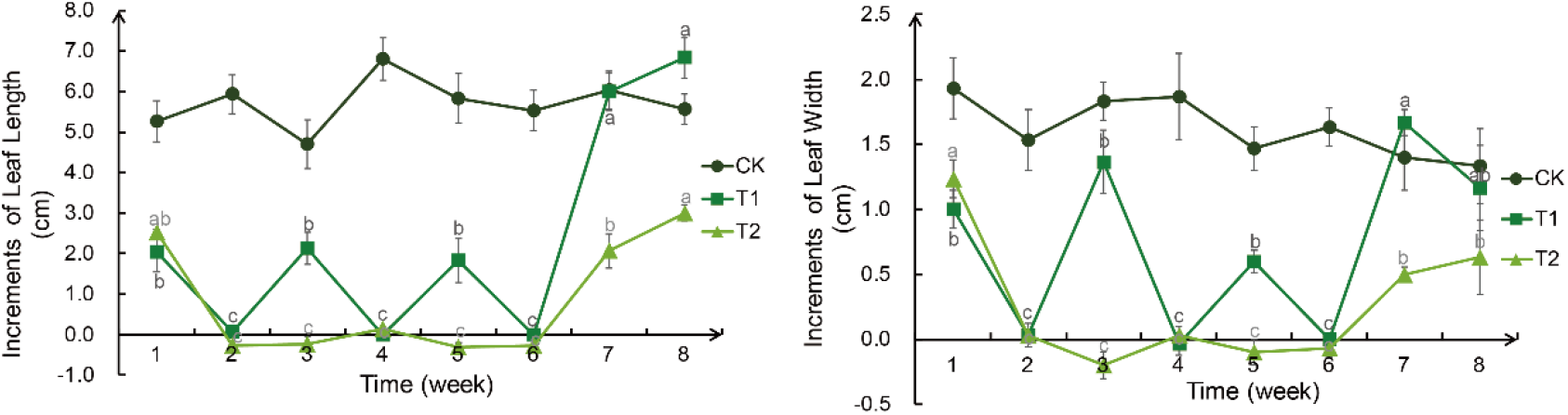
Weekly changes in increments of leaf length and leaf width of the bird’s nest fern under different degrees of drought treatment. a: increments of leaf length, b: increments of leaf width. Significance analysis results: different letters represent significant differences, the same letters represent not significant differences. Significant level was 0.05.

### Measurement results of physiological parameters

RFWC of bird’s nest fern leaves in CK group has been maintained at about 90%(Figure 2). After drought treatment, the leaf relative water content in T1 group fluctuated between 60% and 80%, and that in T2 group decreased to about 40%(Fig. 3). RFWC could be reverted to the normal range after rehydration for two weeks in T1 and T2.

**Fig. 2.**
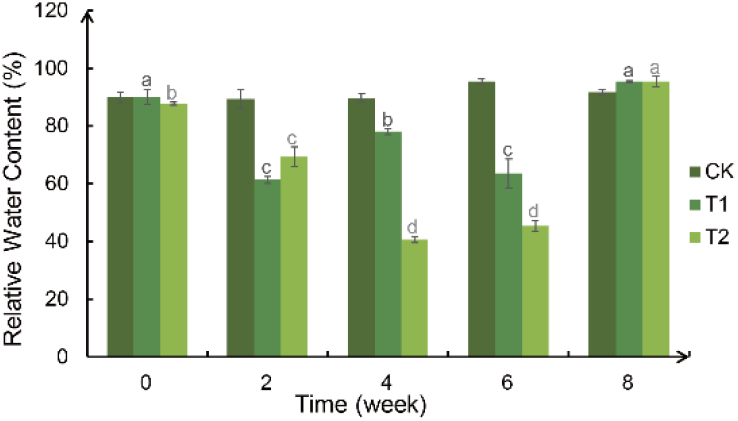
Weekly changes in increments of leaf length and leaf width of the bird’s nest fern under different degrees of drought treatment. Significance analysis results: different letters represent significant differences, the same letters represent not significant differences. Significant level was 0.05.

The change trends of *P*_*max*_,*T*_*r*_ and *g*_*s*_, with different degrees of drought were similar(Fig.4). After drought treatment, the three indexes in T1 and T2 groups decreased to near zero, only in T1 group increased slightly in the 3rd week, and increased rapidly in both groups after rehydration, and the rising range in T1 group was higher than that in T2 group. The change trend of C_i_ was opposite to that of the other three indexes. After drought treatment, the intercellular CO_2_ concentration in T1 group and T2 group was significantly higher than that in CK group, and the rising range in T2 group was larger than that in T2 group. After rehydration, the intercellular CO_2_ concentration in both groups decreased to normal value immediately.

**Fig. 4.**
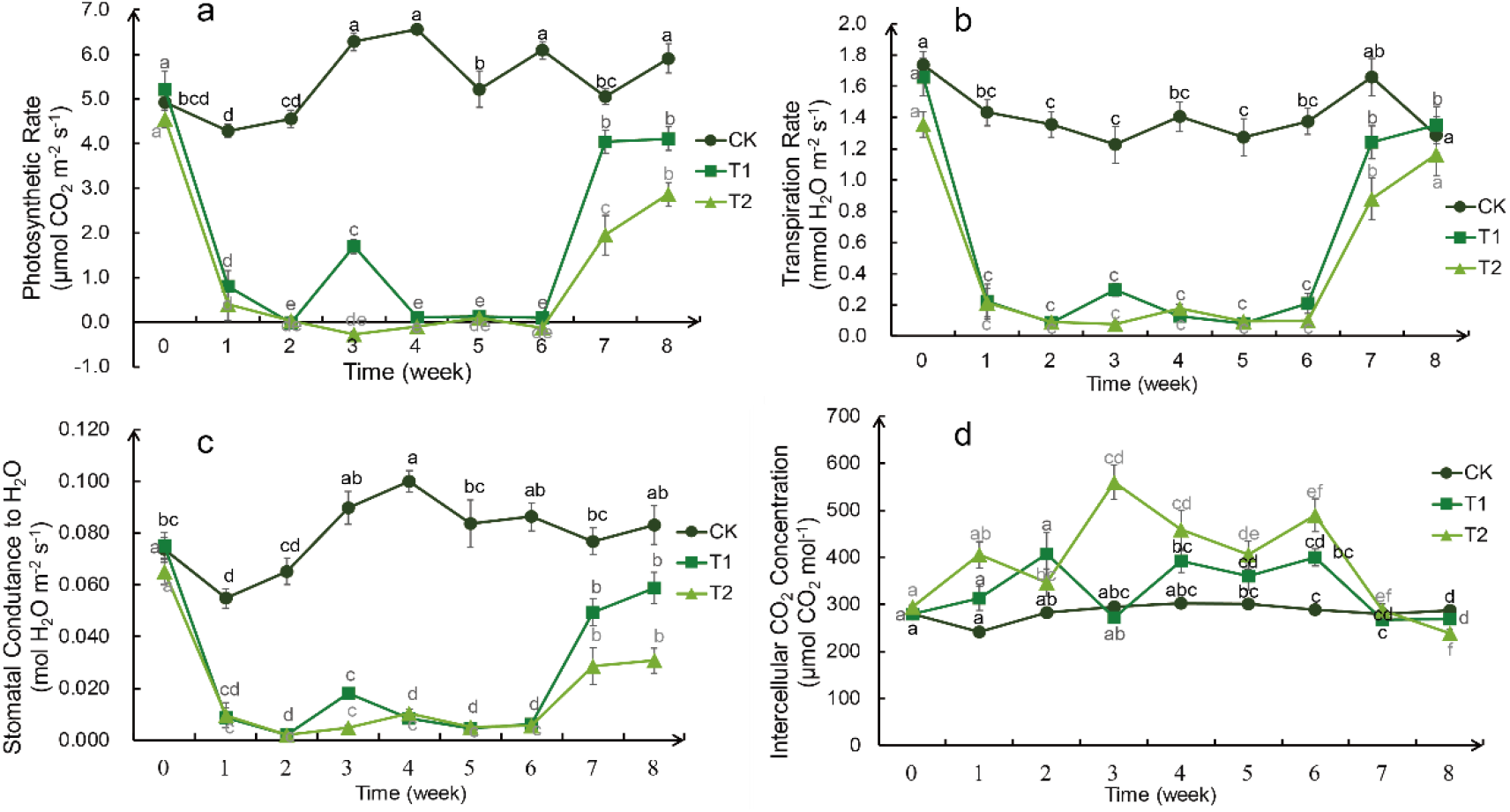
Weekly changes in photosynthetic indicators of the bird’s nest fern under different degrees of drought treatment. a: the maximum net photosynthetic rate (*P*_*max*_), b: transpiration rate (*T*_*r*_), c: stomatal conductance (*g*_*s*_), d: intercellular CO_2_ concentration (*C*_*i*_). Significance analysis results: different letters represent significant differences, the same letters represent not significant differences. Significant level was 0.05.

**Fig. 5.**
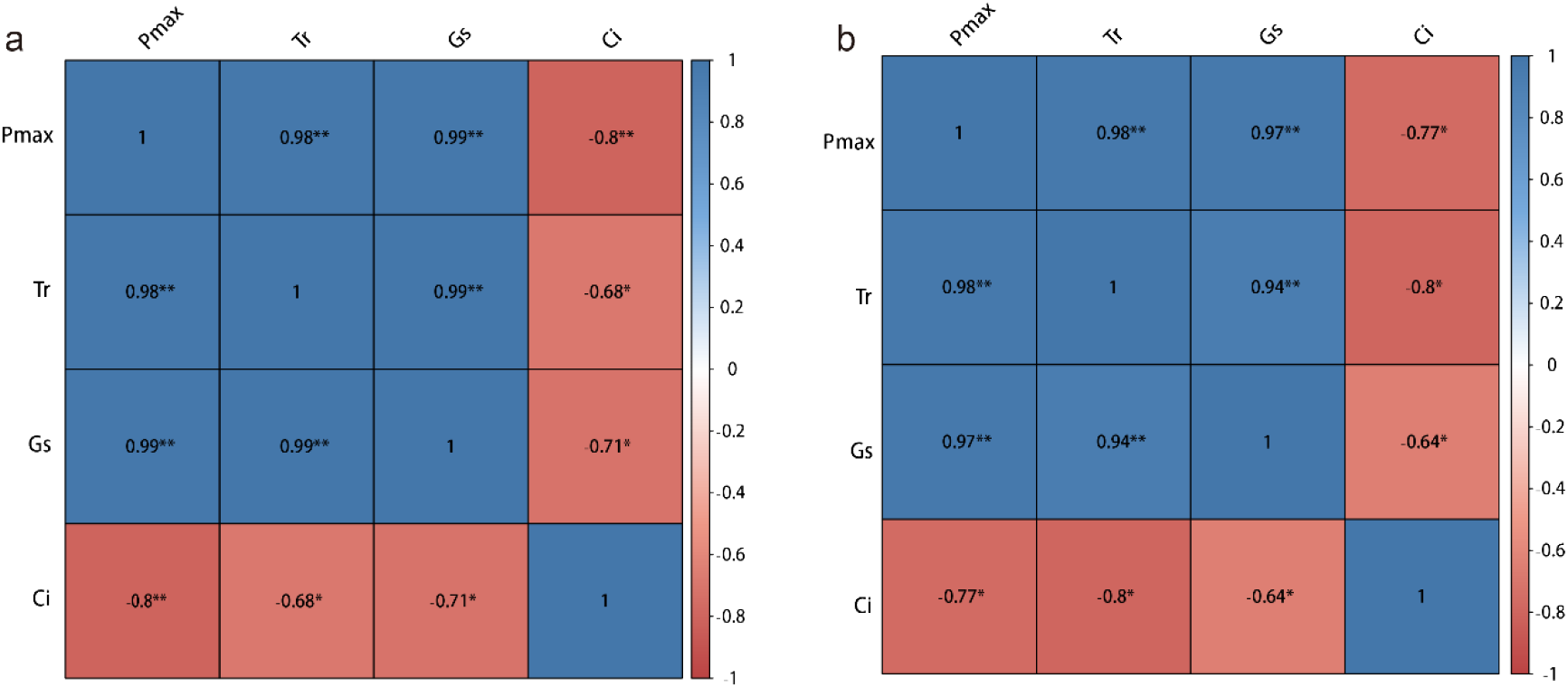
Heat map for correlation analysis between *P*_*max*_, *T*_*r*_, *g*_*s*_ and *C*_*i*_. a: correlation of the T1 group, b: correlation of the T2 group. Where numbers represent relevance, blue represents positive correlation and red represents negative correlation. The darker the color, the greater the correlation.* Stands for significance,* p<0.05 ** p<0.01.

**Fig. 6.**
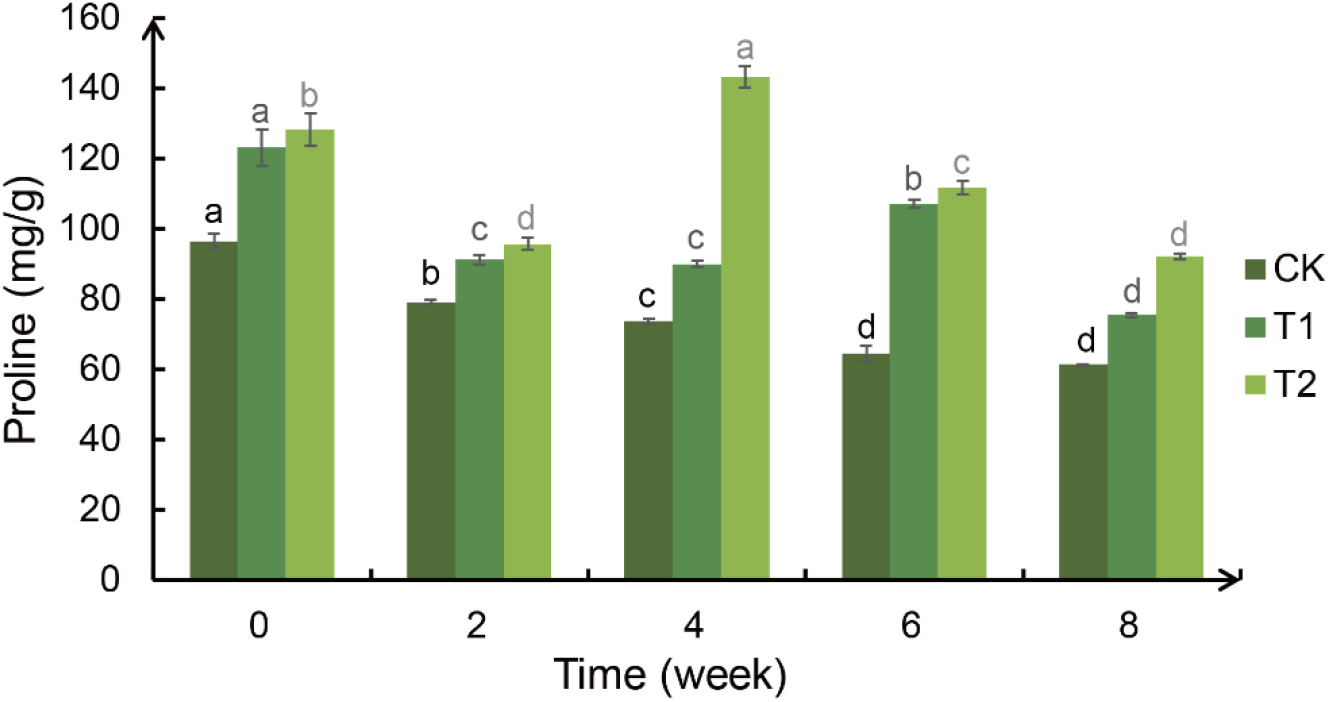
Weekly changes in the proline content of the bird’s nest fern under different degrees of drought treatment. Significance analysis results: different letters represent significant differences, the same letters represent not significant differences. Significant level was 0.05.

We performed a correlation analysis on *P*_*max*_, *T*_*r*_, *g*_*s*_ and *C*_*i*_ of the T1 and T2 groups, respectively. The analysis results showed that there was extremely significant positive correlation between *P*_*max*_, *T*_*r*_, and *g*_*s*_ in the T1 and T2 group (Fig.4), while *C*_*i*_ was negatively correlated with *P*_*max*_, *T*_*r*_, and *g*_*s*_, respectively. In the T1 group, there was extremely significant negative correlation between *C*_*i*_ and *P*_*max*_, but *C*_*i*_ was significantly negatively correlated with *T*_*r*_ and *g*_*s*_. However, in the T2 group, *C*_*i*_ was significantly negatively correlated with *P*_*max*_, *T*_*r*_ and *g*_*s*_.

### Measurement results of biochemical parameters

The phenomenon of T2> T1> CK appeared at the 0th week of proline content, which may be an experimental error caused by the difference in sampling time between the three groups caused by the sampling sequence when sampling. The proline content in the leaves of the bird’s nest fern in the T1 and T2 groups both decreased slightly and then increased significantly after drought treatment. After rehydration, it dropped to the normal value. The proline content of the leaves of the bird’s nest fern in the T1 group increased significantly at the 6th week of drought treatment, while the T2 group increased significantly at the 4th week of drought treatment.

Combining Fig.7 and Table 1, we can know that the betaine content of the bird’s nest fern leaves will not be significantly different under different drought treatments and different periods, so it can be speculated that the drought treatment will not affect the betaine content of the bird’s nest fern leaves.

**Fig. 7.**
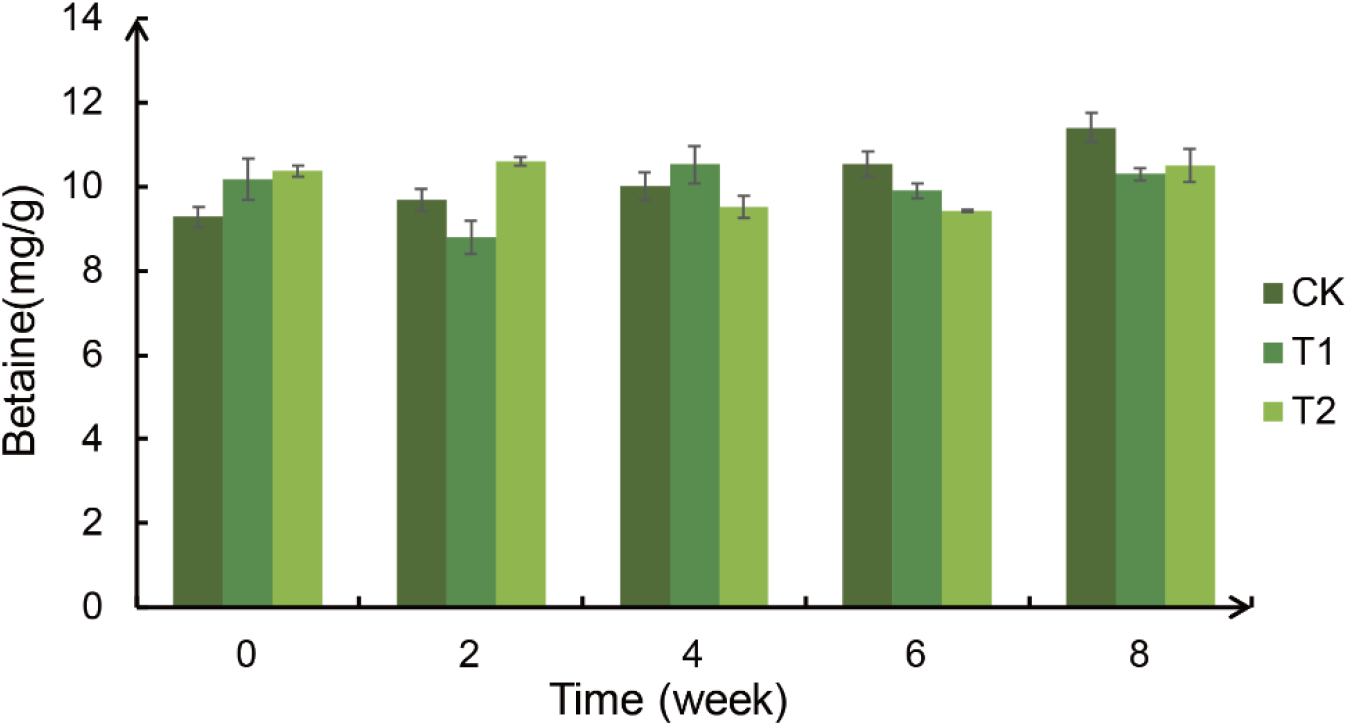
Weekly changes in the betaine content of the bird’s nest fern under different degrees of drought treatment.

**Table 1.** ANOVA analysis of betaine content in leaves of bird’s nest fern treated with different degrees of drought. R^2^: 0.228, * p<0.05 ** p<0.01

| Difference source | Sum of squares | df | Mean square | F | P |
| --- | --- | --- | --- | --- | --- |
| Treatment | 0.449 | 2 | 0.224 | 0.423 | 0.658 |
| Time | 5.494 | 4 | 1.374 | 2.59 | 0.052 |
| Residual | 20.155 | 38 | 0.53 |  |  |

**Table 2.**
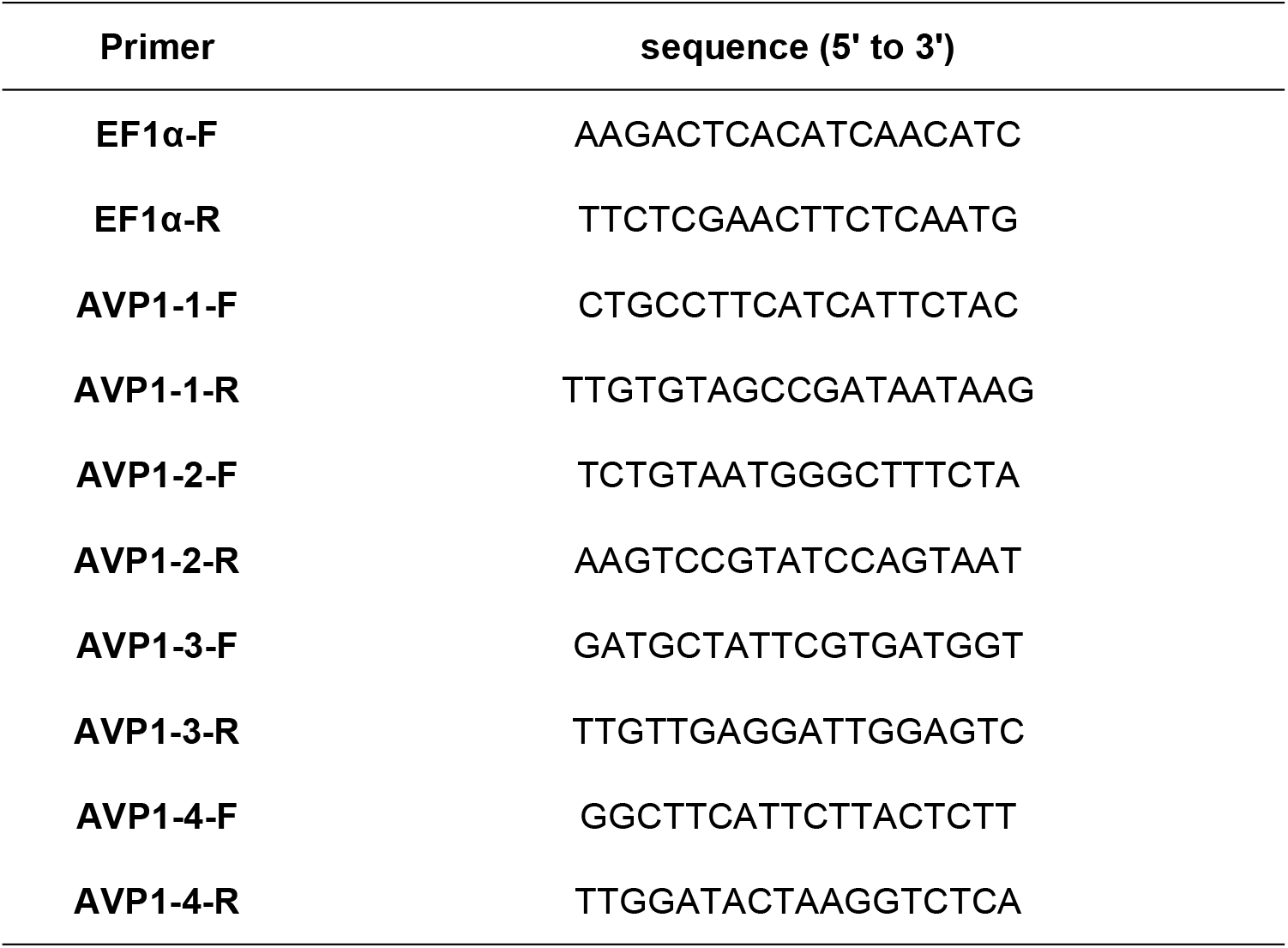
The primers of RT-qPCR. EF1**α**: reference gene, F: forward primer, R: reverse primer

The SOD enzyme activity in the T1 group increased slightly after drought treatment and did not decrease after rehydration, while the T2 group increased significantly during the 2nd week of drought treatment, then gradually decreased in the later period, and decreased to the normal range after rehydration(Fig.8). The POD enzyme activity in the T1 group increased slightly in the 2nd and 6th week, while the T2 group increased significantly in the 4th and 6th week, and they all returned to the normal range after rehydration. CK, T1, and T2 had the same trend of CAT enzyme activity but T2> T1> CK as a whole. All the three treatments increased significantly twice in the 4th and 8th week (the 2nd week after rehydration).

**Fig. 8.**
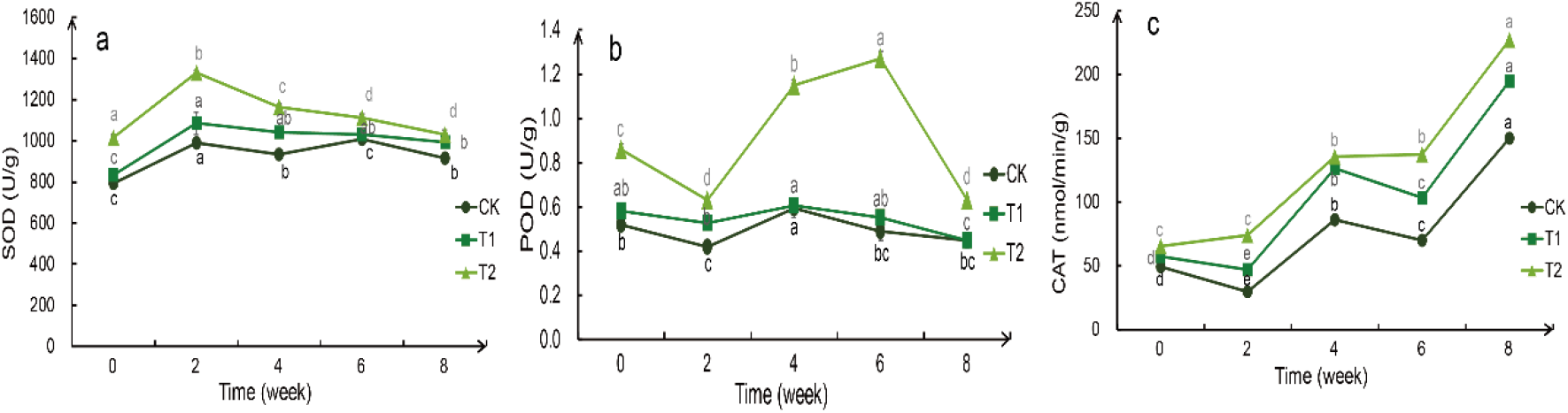
Weekly changes in the enzyme activity of the bird’s nest fern under different degrees of drought treatment.. a: enzyme activity of SOD, b: enzyme activity of POD(*T*_*r*_), c: enzyme activity of CAT. Significance analysis results: different letters represent significant differences, the same letters represent not significant differences. Significant level was 0.05.

### Bioinformatics and gene expression analysis

We identified the conserved domain by the sequence of the gene protein with identified function. The conserved domain of the known *AVP1* gene (AT3G55360.1) was obtained by searching the protein sequence of the known *AVP1* gene (AT3G55360.1) in the PfamA database, and the number PF03030.15, was named H_PPase in the PfamA database. was named H_PPase in the PfamA database. This domain accounts for more than 95% of the total *AVP1* protein sequence. The total length of the protein sequence of *AVP1* gene is 770 amino acids, and its conserved domain is from 21th amino acids to 755th amino acids (Fig. 9).

**Fig. 9.**
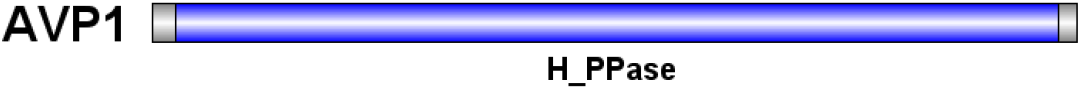
Domains of *AVP1* protein. The gray part is the full length of the protein sequence, and the blue part is the position of the conserved domain

A total of 16 members of proton pyrophosphatase (*AVP1*) gene family were identified from the protein model database of bird’s nest fern, which contained the domain numbered PF03030.15. By observing the phylogenetic tree(Fig.10), it was found that the AVP1 gene was divided into two subfamilies: *AVP* and *AVPL*, in the model plant *Arabidopsis thaliana*, with three sequences *AVP1, AVPL1* and *AVPL2*, selected the bird’s nest fern sequence which was closest to these three sequences and named it *AVP1-1,AVP1-2,AVP1-3,AVP1-4* respectively.

**Fig. 10.**
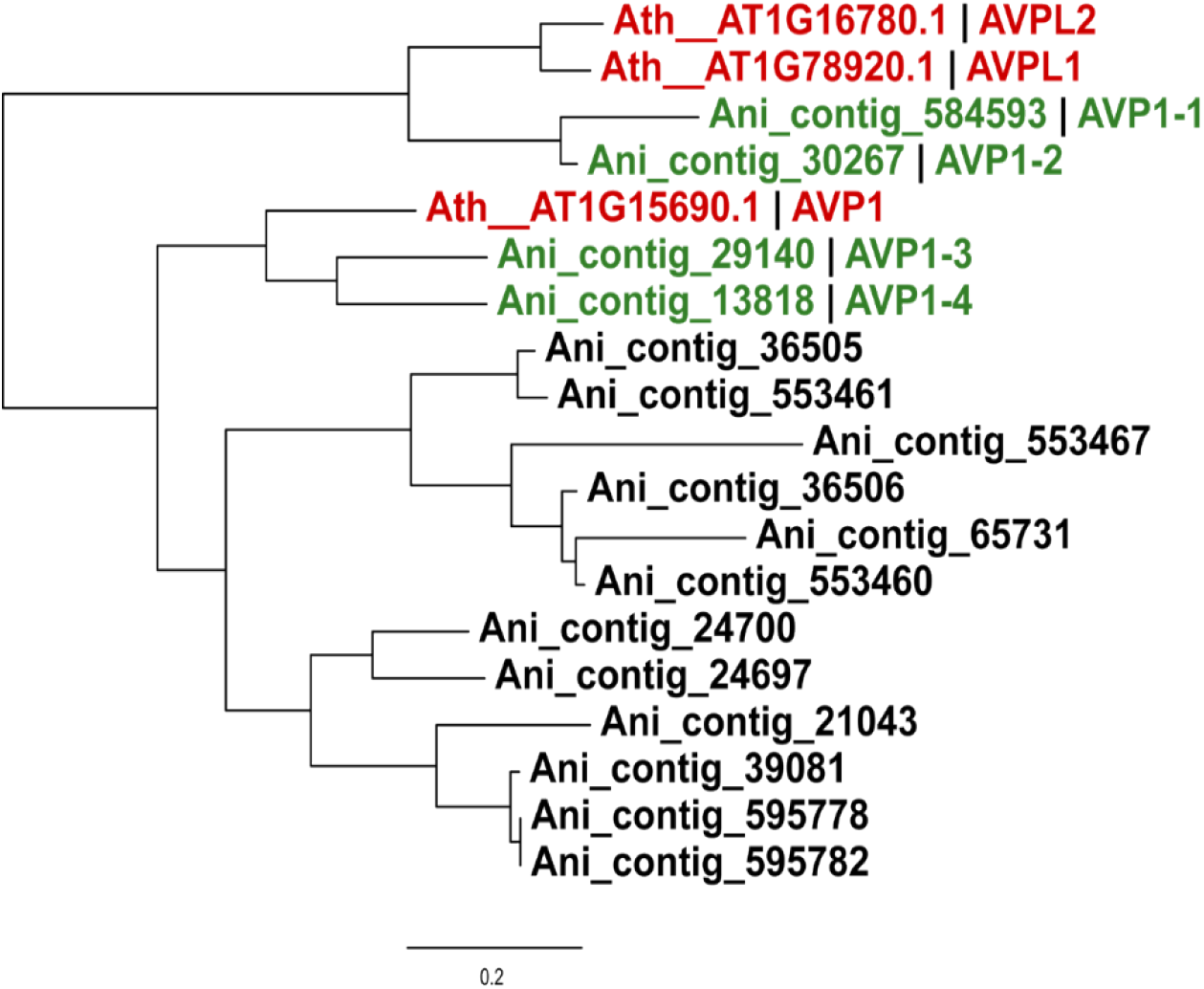
Phylogenetic tree for *AVP1* family members in *Asplenium nidus*. Ath is the abbreviation of *Arabidopsis thaliana*, and Ani is the abbreviation of *Asplenium nidus*.

The primers of RT-qPCR were designed by using Primer5 and Oligo7 software in the Table2.Using EF1a as an internal reference gene, the gene expression of *AVP1-1, AVP1-2, AVP1-3*, and *AVP1-4* were analyzed. It is worth noting that compared with CK, the relative expression of *AVP1-2* in T1 group increased significantly in the 6th week, while that in T2 group increased significantly in the 2nd and 4th week, and decreased in both groups after rehydration(Fig.11). Compared with CK, the relative expression of *AVP1-4* in T2 group increased significantly in the 4th week. The expression of *AVP1-1* and *AVP1-3* was not related to the degree of drought and the time of drought.

**Fig. 11.**
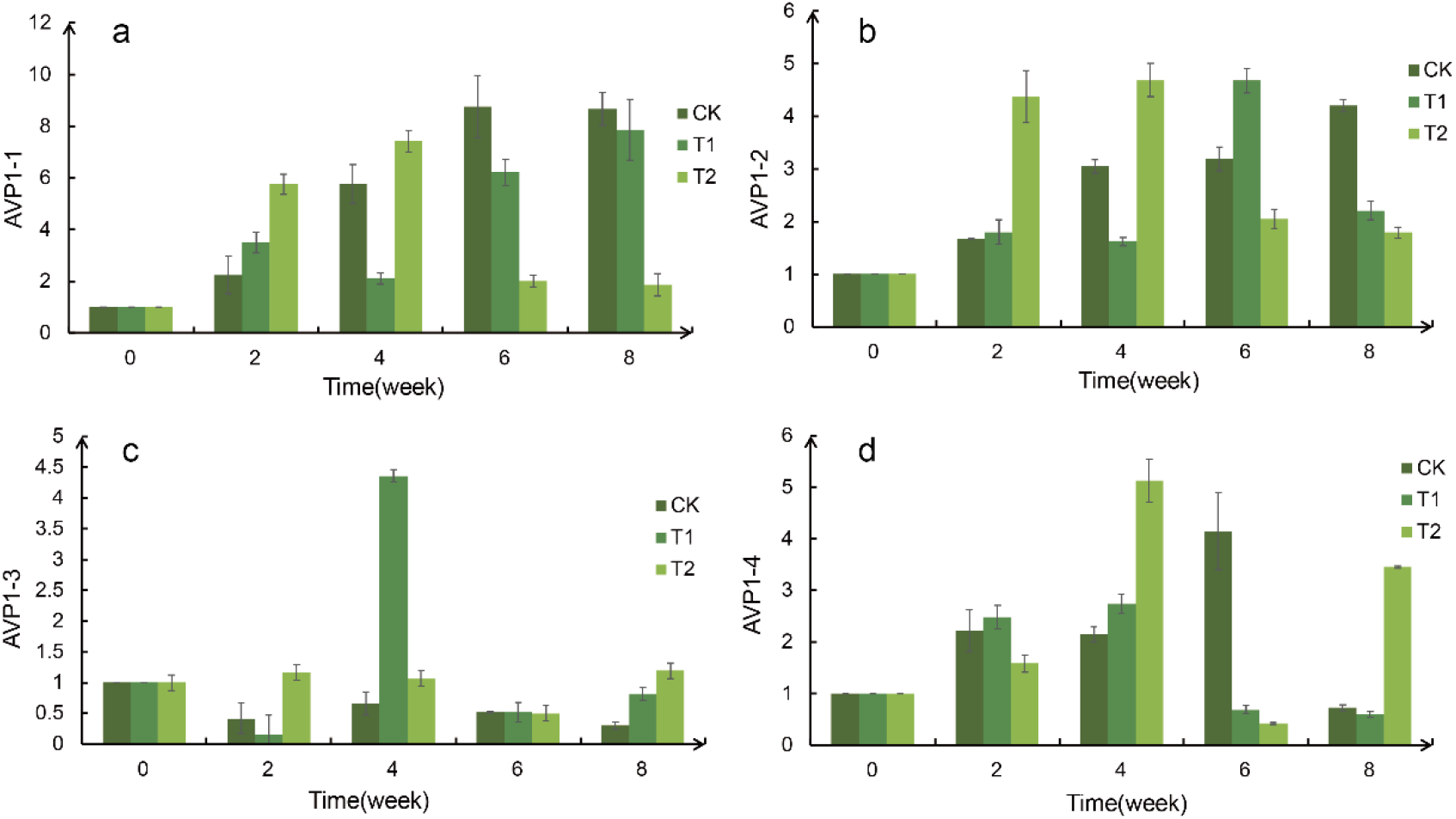
Relative expression of *AVP1-1, AVP1-2, AVP1-3*, and *AVP1-4*. a:*AVP1-1*,b:*AVP1-2*,c: *AVP1-3*,d:*AVP1-4*.

## DISCUSSION

Leaf is an important organ of plant, affecting plant photosynthetic capacity and ornamental value. The growth of leaves is usually inhibited by drought stress. The results of this experiment showed that the leaves of *Asplenium nidus* stopped growing immediately under drought stress, but could recover immediately after rehydration. Even without watering for 6 weeks, the growth rate can gradually return to the normal level with the increase of rewatering time.

The RFWC is an important index to study the drought tolerance of plants. Drought-tolerant plants generally show that they can maintain a relatively high water content even under drought stress, or that they can recover quickly after rehydration, although they are reduced under drought stress.(Wang et al., 2019) Although it decreases greatly under drought stress, it can recover quickly after rehydration. The experimental results show that although the RFWC of *Asplenium nidus* decreased greatly under drought stress, which can be reduced from 90% to 40% without watering at all, but rehydration can help it quickly return to the normal value after 6 weeks without watering. The results are consistent with to ZHANG’s (Q. ZHANG et al., 2009)study of two kinds of epiphytic pteridophytes, which are *Neottopteris nidus* (L.) and *Microsorium punctatum*. This experiment showed that although RFWC of *Asplenium nidus* decreased greatly after drought stress, it could remain above the harmful water content, recover immediately after rehydration. As a result, the *Asplenium nidus* has strong adaptability to the change of environmental water content.

The photosynthetic characteristics of plant leaves have certain adaptability to the habitat, and some plants will reduce photosynthesis to adapt to water deficit. This study showed that the P_max_, T_r_ and g_s_ of *Asplenium nidus* leaves decreased significantly to about 0 under drought stress, and recovered quickly after rehydration. These three indexes were significantly positively correlated with each other, while the intercellular CO_2_ concentration was significantly negatively correlated with them. This phenomenon shows that bird’s nest fern leaves limit the utilization of light energy and CO_2_ by limiting stomatal opening. To inhibit the efficiency of plant photosynthesis to adapt to drought, which is supported by K. NISHIDA et al(K. NISHIDA et al., 2017).

Osmotic regulation is an important way for plants to maintain normal metabolic activities by changing intracellular solute concentration and resulting in changes in cell osmotic potential under drought stress. In this study, proline and betaine, two recognized osmotic regulators, were determined. The results showed that the proline content of bird’s nest fern leaves was not sensitive to drought stress, but increased in the later stage of drought stress, which was consistent with the results of Huang Yong’s study(Huang Yong, 2012) on *Neottopte nidus*. The results showed that the regulation of proline in the leaves of bird’s nest fern would not be started until the drought stress reached a certain degree, but the content of betaine did not change with the degree and time of drought. This result is consistent with the findings of Fallard,A et al (Fallard et al., 2018)in *Hymenoglossum cruentum* and *Hymenophyllum dentatum*. Fallard,A et al believe that the conventional solute accumulation in higher plants is not fully adapted to the drought tolerance mechanism of pteridophytes, which is verified by our study.

However, when the plant is under water stress, the accumulation of reactive oxygen species is accelerated. When the concentration reaches a certain threshold, the macromolecular substances in the plant cells will undergo peroxidation, which can affect the normal growth of the plant. Some plants have formed a complete active oxygen scavenging system during a long-term evolution. SOD catalyzes the O_2_^-^· disproportionation reaction to H_2_O_2_ and O_2_. POD and CAT are mainly responsible for the H2O2 produced by the disproportionation reaction. It is generally considered that during drought stress, the changes in protective enzyme activity of drought-resistant plants first rise and then decrease, while others show a constant or slightly increased trend. The enzyme activities of non-drought-resistant plants appear as continued decline, SOD and POD in the leaves of the bird’s nest fern under drought stress are in line with this trend in this study.

In summary, this study indicates that *Asplenium nidus* stops the growth of the stomata by limiting the opening of the stomata, thereby limiting the use of light energy and CO_2_, inhibiting plant photosynthesis and reducing life activities to adapt to drought. And in order to resist drought stress, the *Asplenium nidus* changes the osmotic potential by increasing the proline content to maintain normal metabolic activities and prevents the damage of active oxygen by increasing the enzyme activities of SOD and POD. and changing the penetration by increasing the proline content. It is expected to maintain normal metabolic activities under stress conditions and prevent the damage of reactive oxygen species to resist drought by increasing the enzyme activities of SOD and POD. *Asplenium nidus* is a plant with drought tolerance.

A large number of studies have shown that *AVP1* plays an indispensable role in important physiological processes such as plant growth and stress resistance. (Li et al., 2005; Qin et al., 2012)Therefore, in this study, the *AVP1* gene in Bird’s Nest Fern was selected for gene identification and expression analysis. *AVP1-1, AVP1-2, AVP1-3, AVP1-4* were screened through three steps: identification of conserved domains, mining of protein sequences containing target domains, and construction of phylogenetic trees containing sequences of target domains *AVP1* family genes in two bird’s nest ferns. The expression of these four genes under drought stress was analyzed by RT-qPCR. This study speculates that *AVP1-2* and *AVP1-4* may be drought-resistant genes of *Asplenium nidus*. Drought resistance research in *Asplenium nidus* is still in its infancy. Only a few articles have studied the effects of drought stress on it, and few studies have focused on drought tolerance genes. Therefore, we believe that in-depth research on *Asplenium nidus*. should be carried out in the next stage in order to establish a complete drought-resistant system of *Asplenium nidus*.

## Acknowledgement

This work was financially supported by grants from National Natural Science Foundation of China (NSFC No.31660229).

